# Physiologically Relevant Coculture Model for Oral Microbial-Host Interactions

**DOI:** 10.1101/2025.01.10.632380

**Authors:** Zeyang Pang, Nicole Cady, Lujia Cen, Thomas M. Schmidt, Xuesong He, Jiahe Li

## Abstract

Understanding microbial-host interactions in the oral cavity is essential for elucidating oral disease pathogenesis and its systemic implications. *In vitro* bacteria-host cell coculture models have enabled fundamental studies to characterize bacterial infection and host responses in a reductionist yet reproducible manner. However, existing *in vitro* coculture models fail to replicate the physiological oxygen gradients critical for studying these interactions. Here, we present an asymmetric gas coculture system that simulates the oral microenvironment by maintaining distinct normoxic and anaerobic conditions for gingival epithelial cells and anaerobic bacteria, respectively. Using *Fusobacterium nucleatum*, a key oral pathobiont, we demonstrate that the system preserves bacterial viability and supports the integrity of telomerase-immortalized gingival keratinocytes. Compared to conventional models, this system enhanced bacterial invasion, elevated intracellular bacterial loads, and elicited more robust host pro-inflammatory responses, including increased secretion of CXCL10, IL-6, and IL-8. Additionally, the model enabled precise evaluation of antibiotic efficacy against intracellular pathogens. These results underscore the utility of this coculture platform for studying oral microbial pathogenesis and screening therapeutics, offering a physiologically relevant approach to advance oral and systemic health research.

**TOC:** 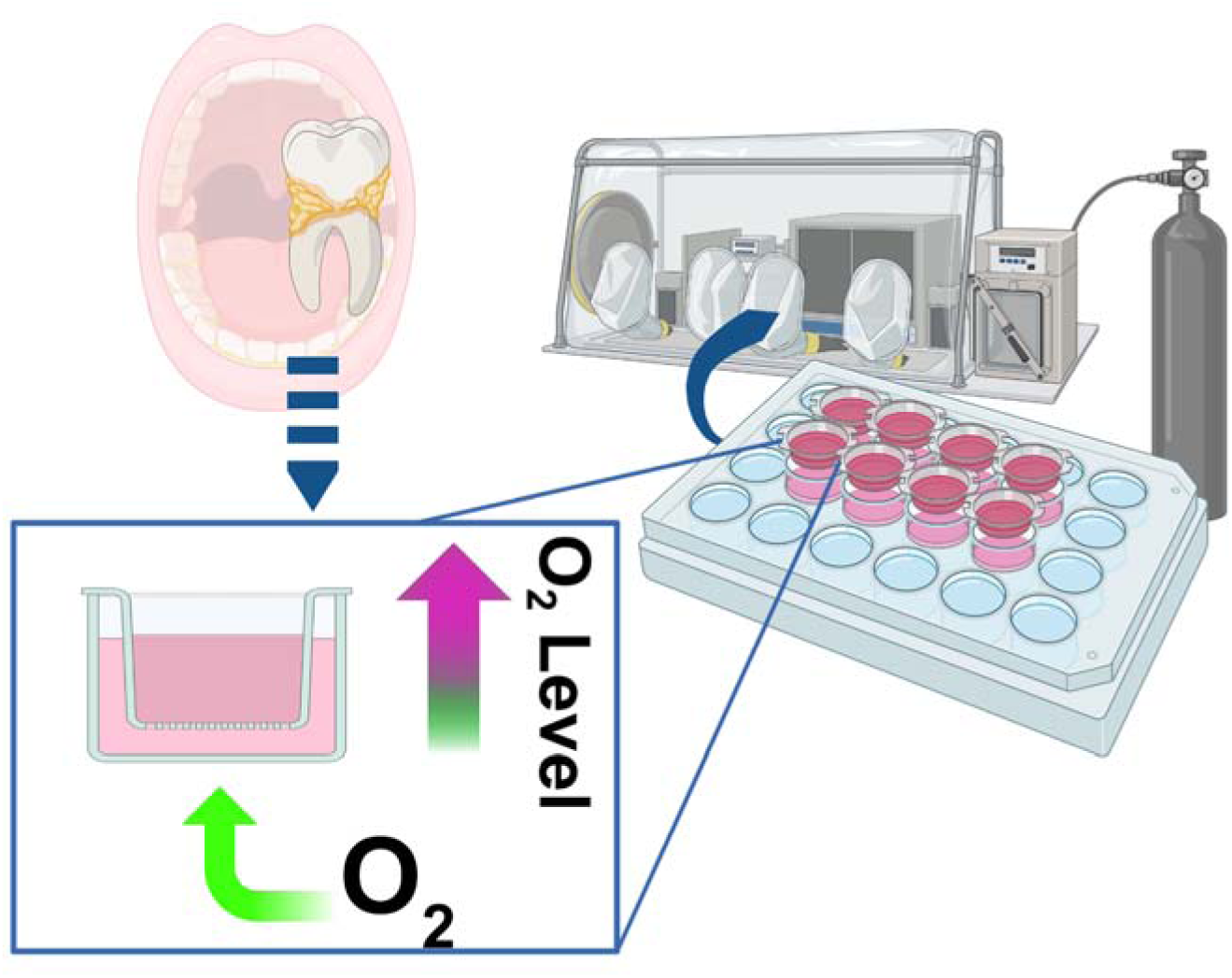

## Introduction

The oral cavity is a complex ecosystem that hosts different species of microorganisms (Attar, 2016). Oral microbes play a crucial role in the development of oral chronic inflammation diseases such as gingivitis and periodontal disease (Signat et al., 2011) (Mei et al., 2020). It has become increasingly appreciated that microbial-host cell interaction affects local immune responses in the oral cavity and may also profoundly affect systemic health through inflammatory pathways (Gao et al., 2018) (Le Bars et al., 2017). To maintain homeostasis, the gingival epithelium provides a physical and biological barrier responsible for host defense yet is frequently challenged by pathogenic bacteria. Therefore, understanding the interaction between oral bacteria and host tissues is essential for both local oral health and the prevention of systemic diseases. To understand the exact role of oral microorganisms in diseases, the coculture system, which allows for the simultaneous cultivation of bacteria and host cells, has become an essential tool in this field. By simulating the interaction between anaerobes and host cells, we can better understand the pathogenic mechanisms of these microorganisms in certain disease models and even explore their potential role in systemic diseases (Fofanova et al., 2019). Therefore, studying the coculture of oral microorganisms and host cells is significant in promoting a scientific understanding of the relationship between the oral cavity and systemic health.

However, there are significant limitations in current bacterial-host cell coculture systems, particularly when involving anaerobic bacteria (Zhang et al., 2021). In traditional cell culture systems, anaerobic bacteria are typically introduced directly into the normoxic culture medium, which presents several challenges. First, anaerobic bacteria often struggle to maintain viability in these environments due to the presence of oxygen (Jalili-Firoozinezhad et al., 2019). This significantly impairs their growth and metabolism, limiting the duration of studies to less than 12 hours (Mountcastle et al., 2020). Second, the oral environment naturally contains distinct normoxic and anaerobic gradients, exposing cells and bacteria to varying oxygen levels (Xu et al., 2015). Without replicating these oxygen gradients, it is difficult to fully simulate the physical and chemical properties of the oral cavity, making coculture systems less representative of actual physiological conditions (Albenberg et al., 2014). Lastly, the conventional method fails to effectively simulate the complex process by which bacteria invade host cells, limiting the relevance and accuracy of the long-term models in reflecting *in vivo* conditions (Mountcastle et al., 2020). As a result, these limitations hinder the ability to fully understand oral bacterial pathogenesis and necessitate the development of improved coculture techniques that better simulate the natural interactions between anaerobic bacteria and host cells. To date, *in vitro* devices to mimic the gingival pockets are largely complex and do not necessarily allow for facile characterization of microbe-host interactions (Makkar et al., 2023) (Adelfio et al., 2023).

To address these gaps, we propose an asymmetric gas coculture system to recapitulate an oxygen gradient within the interface of the host and oral anaerobes, which allows for studying bacterial invasion and infection under physiologically relevant conditions (**Figure 1A**). Although this setup has been employed for interrogating the gut microbe-host interactions *in vitro*, to the best of our knowledge, a similar system has not been used for characterizing the interplay between oral microbes and the host (Fofanova et al., 2019). As a proof-of-concept, we employed *Fusobacterium nucleatum* (*Fn*), a key oral pathobiont, to demonstrate the utility and advantages of our system. *Fn* is of particular interest due to its ability to invade gingival epithelial cells and its association with both periodontal disease and systemic conditions such as gastrointestinal cancers (Lai et al., 2022) (Stokowa-Sołtys et al., 2021). We demonstrate that our coculture system outperforms traditional platforms by recapitulating the oxygen gradient, allowing oral anaerobes to thrive under anaerobic conditions while maintaining oral epithelial cells at physiological oxygen levels (i.e., the normoxic condition). This approach holds the potential to advance our understanding of microbial pathogenesis in oral and systemic diseases and serves as a valuable platform for drug screening and therapeutic development.

**Figure 1:**
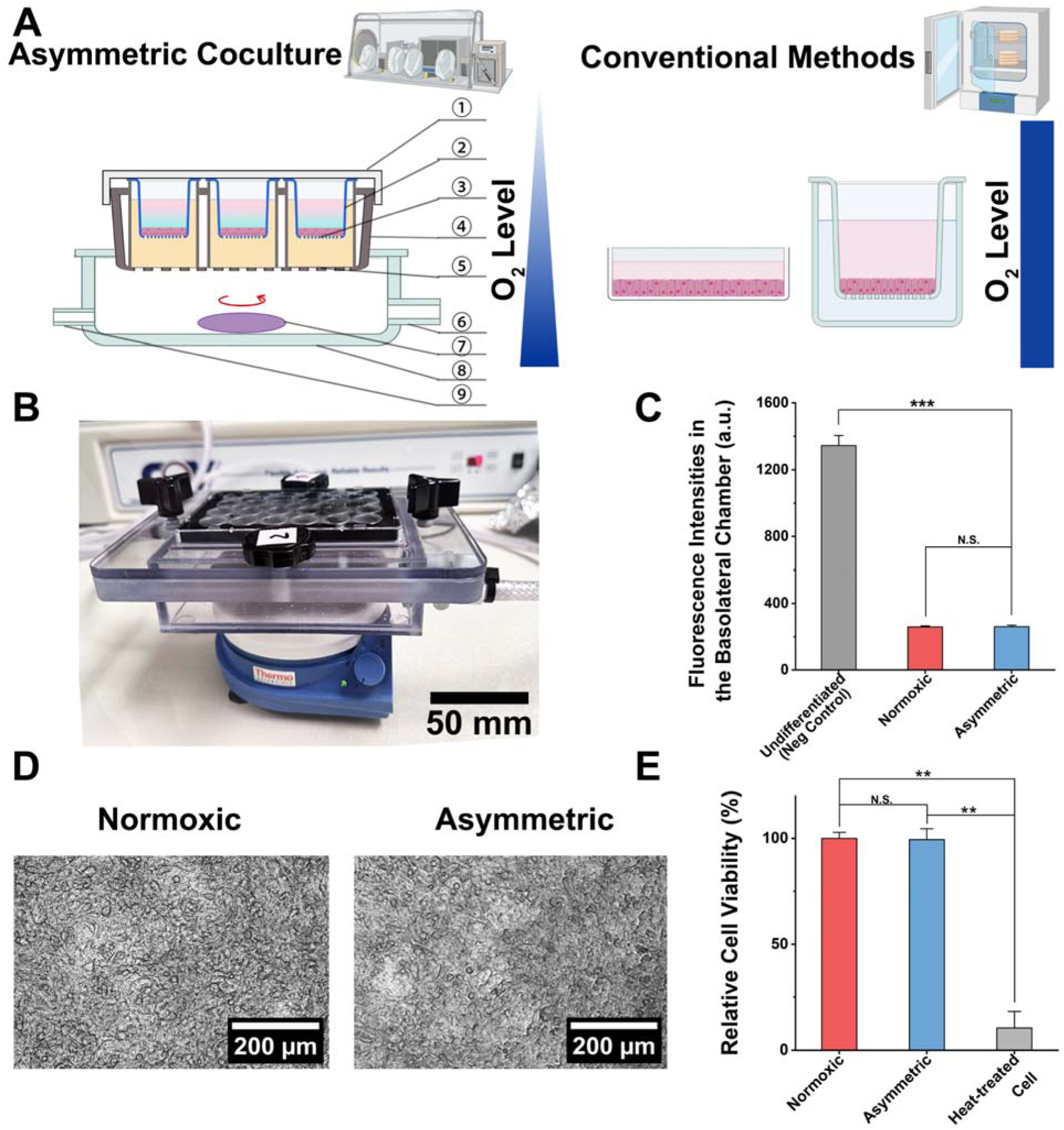
Development of an asymmetric gas coculture system utilizing an immortalized gingival epithelial cell line as a barrier. (A) A schematic comparing the asymmetric coculture chamber with a ventilation panel to the traditional methods for mono- and coculture of oral gingival epithelial cells. The plate cap ① covers the gas-permeable plate ④, preventing contamination and leakage. The apical chamber of the Transwell inserts ② is filled with cell-compatible media (CCM, see **Table S1**), which supports the stable growth of the cell monolayer ③ and *Fusobacteria*. The lower compartment (i.e., the basolateral compartment) of the Transwell insert is filled with cell culture medium. The gas-permeable membrane ⑤ at the bottom ensures unidirectional oxygen diffusion, creating an oxygen gradient throughout the coculture chamber. Basolateral gas flow containing 10% oxygen enters through the gas inlet ⑨, spreading evenly through the asymmetric coculture chamber ⑧ with a magnetic stirrer ⑦. Exhaust gas is discharged through the gas outlet ⑥, completing the system’s airflow (Fofanova et al., 2019). The figure was created using BioRender. (B) A physical picture of the asymmetric coculture chamber. (C) Comparison of fluorescence intensities of FITC-dextran in the basolateral chamber at 24 h after adding FITC-dextran to the apical chamber of Transwells containing TIGK monolayers. Undifferentiated (negative control) and differentiated TIGKs under the normoxic culture conditions (referred to as “normoxic”) were compared to differentiated TIGKs under the asymmetric culture condition (referred to as “asymmetric”). The background fluorescence intensity of the blank culture medium was subtracted for each condition. Undifferentiated TIGK monolayer cultured under the normoxic condition before switching to a differentiation medium containing Ca^2+^ serves as the negative control. (N.S.: *p*>0.05, ***: *p*<0.001, n=2 technical replicates, N=3 biological repeats). (D) The morphology of TIGK monolayers in Transwell inserts maintained in a cell culture incubator under normoxic conditions or cultured in an asymmetric coculture chamber for 24 h. The collagen coating is known to affect bright field imaging due to the presence of the collagen fibers, which can obscure delicate details of the cells or structures being observed compared to an uncoated surface (Hashimoto et al., 2020). (E) Comparison of cell viability in TIGK monolayers cultured under normoxic and asymmetric culture conditions. Heat-treated cells are the negative control (N.S.: *p*>0.05, **: *p*<0.01, n=3, N=3).

## Results

### The asymmetric gas chamber maintains the viability of telomerase-immortalized gingival keratinocytes (TIGKs)

Different regions in the oral cavity are exposed to varying levels of oxygen, influenced by factors such as proximity to oxygen-rich surfaces (like the open air and saliva) and oxygen-consuming bacterial communities. At the bacterial-host interface, these gradients become even more pronounced due to the metabolic activity of both the host cells and the microbial community (Kenney and Ash, 1969) (Mettraux et al., 1984). Thus, an asymmetric gas chamber was developed to mimic the distinct oxygen gradients in the oral cavity. The experimental setup utilized a monolayer of telomerase-immortalized gingival keratinocytes (TIGKs) cultured on Transwell inserts (**Figure 1A**). The apical compartment was maintained under anaerobic conditions, while the basolateral chamber was supplied with a 10% oxygen gas mixture, simulating two distinct culture environments (**Figure 1A**). This setup contrasts with conventional methods, which expose host cells and anaerobes to uniform normoxic conditions. To recapitulate an oxygen concentration gradient within the interface of oral epithelial cells and anaerobes established by the gingival epithelium, we used the TIGK cell line, which was previously developed to enable *in vitro* studies while maintaining biological characteristics similar to human primary gingival epithelial cells (Bailey et al., 2013).

Before transferring TIGKs into the asymmetric gas coculture chamber, the cells—exhibiting a polygonal shape, close packing, and large, round nuclei—were first seeded in Transwell inserts in Dermal Cell Basal Medium (**Figure S1**) (Holm and Qu, 2022). After 24 h of attachment, the medium was replaced with Dulbecco’s Modified Eagle Medium (DMEM), in which calcium ions promoted differentiation of the TIGKs, leading to the formation of a tight monolayer on the collagen IV-coated membrane in Transwells (Leonardo et al., 2020). Following 4 d of differentiation, the intact monolayer was examined under a microscope (**Figure S2**). After confirming monolayer formation, we transferred the Transwell inserts containing TIGK monolayers to the asymmetric gas coculture chamber and placed the entire setup in an anaerobic workstation (**Figure 1B**). Unlike typical Transwell assays where both apical and basolateral chambers utilize mammalian cell culture media to support the growth of host cells, we developed a chemically defined medium (cell-compatible medium, CCM) introduced in the apical chamber to allow for the maintenance of anaerobes without affecting the viability of TIGKs in the coculture. After placing the asymmetric gas chamber in the anaerobic workstation, the basolateral chamber was stabilized for 3 h with a continuous supply of 10% oxygen at 0.5 standard cubic feet per hour (SCFH) to establish an oxygen gradient. This setup contrasts with the conventional methods, where TIGKs were cocultured with oral anaerobes under a uniform normoxic culture condition in a 5% CO_2_ incubator (Kasper et al., 2020) (Casasanta et al., 2020) (Udayasuryan et al., 2022).

We first assessed whether the asymmetric gas chamber could affect the integrity of the TIGK monolayer in the absence of bacterial challenge. We used fluorescein isothiocyanate (FITC)-labeled dextran (molecular weight=4,000 Da) as a small molecule surrogate to characterize the formation of a tight biological barrier established by the differentiated TIGK monolayer and compared it to the undifferentiated state. 24 h after introducing FITC-dextran in the apical chamber, the fluorescence intensity of the media in the basolateral chamber was quantified to indicate the leakiness of the TIGK monolayer (**Figure 1C**). Compared to undifferentiated TIGKs, the diffusion of FITC-dextran through the Transwell under the normoxic condition decreased more than five times, suggesting the formation of a tight monolayer barrier. Moreover, the intactness of the differentiated monolayer under the asymmetric condition remained comparable to that of the normoxic condition. Additionally, bright-field imaging of the monolayers showed minimal changes with regards to cell shape or monolayer intactness after 24 h under asymmetric conditions compared to normoxic conditions (**Figure 1D**). Notably, the collagen coating affects bright field imaging due to the presence of collagen fibers, which can obscure the details of the cells or structures being observed compared to the uncoated surface (Hashimoto et al., 2020) (**Figure S1** and **S2**). The monolayer cells were further dissociated to measure cell viability by flow cytometry. SYTOX Green, a cell viability dye, was applied to quantify the percentage of dead cells. Due to its strong binding affinity to nucleic acids, this dye emits an intense fluorescence in membrane-compromised dead cells. Flow cytometry analysis with SYTOX Green staining indicated no significant increase in cell death under asymmetric conditions compared to normoxic conditions (**Figure 1E**, **Figure S3**). These results validate that the asymmetric gas chamber maintains the viability and functionality of TIGK monolayers, providing a robust platform for the following studies on characterizing the interactions between anaerobic bacteria and gingival epithelial cells.

### The asymmetric gas coculture system preserves the viability of *Fusobacterium nucleatum*

In conventional studies of oral anaerobic bacteria-cell interactions, anaerobes such as *F. nucleatum* (*Fn*) are introduced into well plates or Transwells with epithelial cells or other host cell types in a static incubator containing 5% CO_2_ and ambient oxygen levels (Li et al., 2023) (Koido et al., 2022). We speculate that traditional normoxic conditions can compromise the viability of anaerobes (Han et al., 2000). In comparison, our asymmetric gas coculture chamber places a monolayer of gingival epithelial cells such as TIGKs as a physiologically relevant barrier between the two chambers, allowing anaerobes to be cultured in the apical chamber under anaerobic conditions. This ensures that the monolayer of TIGKs reduces oxygen diffusion from the basolateral chamber to the apical chamber, recapitulating an oxygen gradient present on the mucosal surfaces (Huang et al., 2011). We hypothesized that utilizing this asymmetric coculture chamber can preserve the viability of *Fn* in the apical chamber compared to the conventional coculture methods under normoxic conditions. To validate this hypothesis, we first tested two commonly used *Fn* type strains (*Fn* 23726 and *Fn* 25586). Instead of using a high multiplicity of infection (MOI) in the literature (Liu et al., 2022) (Udayasuryan et al., 2024), which may not be physiologically relevant, we chose an MOI of 1. Bacteria were seeded in the apical chambers of Transwells under asymmetric or normoxic conditions. After 24 h, flow cytometry revealed a significant reduction in bacterial viability under normoxic conditions, with viability decreasing by over 70% for *Fn* 23726. In contrast, the asymmetric gas chamber preserved bacterial viability, comparable to strictly anaerobic conditions (**Figure 2A–C**). Similar findings were observed in *Fn* 25586, showing that the asymmetric gas coculture system could maintain the viability of *Fn* 25586 better than the conventional normoxic coculture condition. We reason that oxygen exposure may have compromised bacterial viability under the normoxic condition, affecting how bacteria interact with gingival epithelial cells (Robinson et al., 2020). In comparison, the asymmetric gas coculture system offers certain advantages when bacterial viability is critical for characterizing anaerobe-host interactions under physiologically relevant conditions.

**Figure 2:**
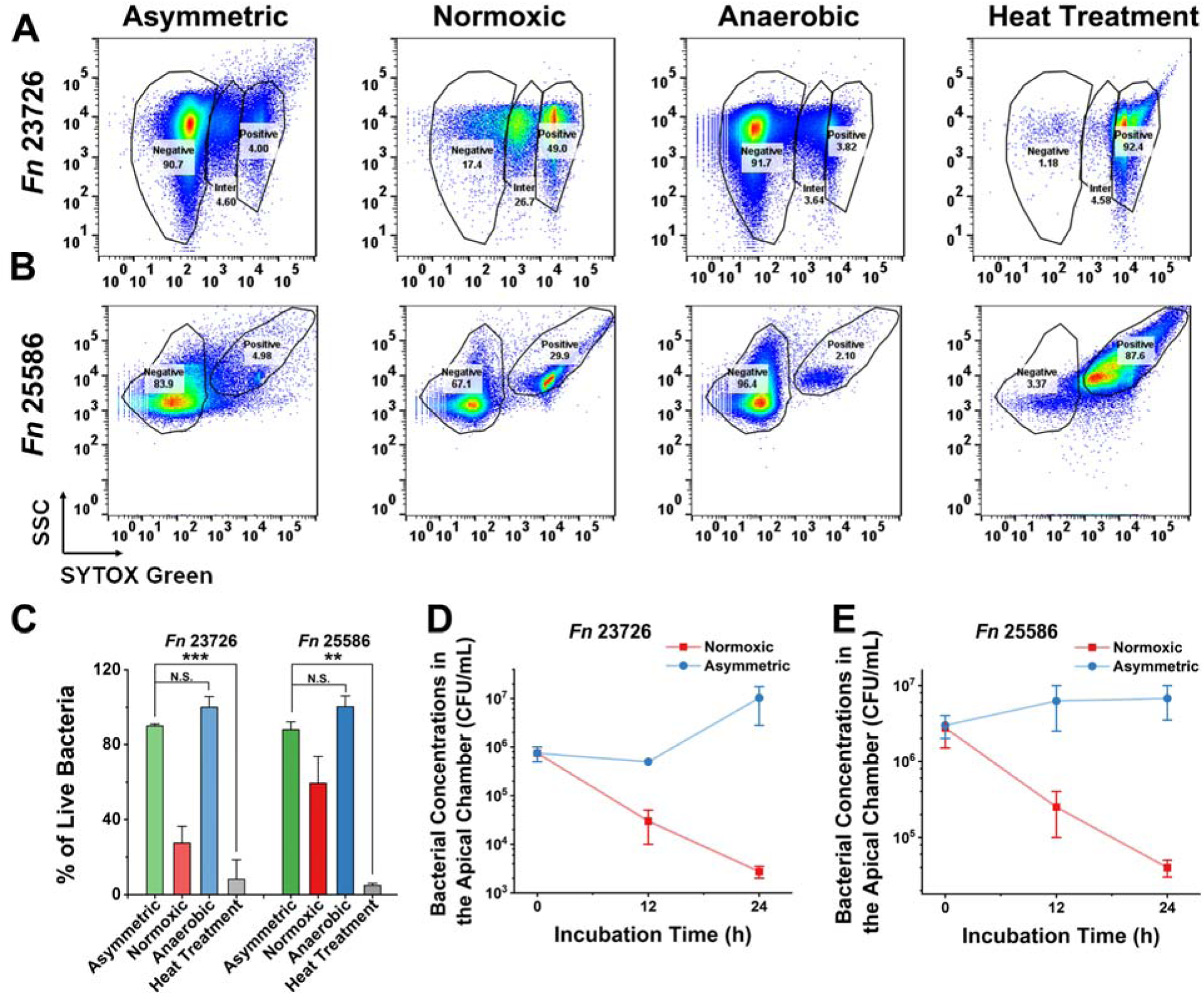
The asymmetric gas setup maintains the viability of *Fn*. Representative flow cytometry results for (A) *Fn* 23726 and (B) *Fn* 25586 under asymmetric gas coculture (with TIGK monolayer), normoxic coculture (with TIGK monolayer), anaerobic condition (in the anaerobic workstation), and dead cell negative control after heat treatment. The MOI was 1. SSC: side scattering. (C) Viability quantification of *Fn* strains 23726/25586 assessed by flow cytometry following different treatment groups, including the asymmetric gas coculture, normoxic coculture, anaerobic culture (positive control), and dead cell negative control after heat treatment (N.S.: *p*>0.05, **: *p*<0.01, ***: *p*<0.001, n=2 technical replicates, N=3 biological replicates). CFUs of (D) *Fn* 23726 and (E) *Fn* 25586 in the apical chamber under normoxic and asymmetric co-culture conditions at 0, 12, and 24h time points (n=2, N=3). The MOI was 1.

In addition to using SYTOX green for measuring bacterial viability, we quantified and compared the temporal changes of colony-forming units (CFUs) of *Fn* 23726 and *Fn* 25586 in the apical chambers of Transwells with differentiated TIGKs under asymmetric or normoxic coculture conditions. At 0, 12, and 24 h post *Fn* inoculation with an MOI of 1, small aliquots were removed from the apical chamber, serially diluted, and plated on Columbia blood agar plates for CFU quantification. For both *Fn* 23726 and *Fn* 25586, relative to the initial bacterial concentrations, there were 10–100-fold decreases in CFUs under the normoxic coculture at the 24-h endpoint, consistent with the reduced viability of *Fn* strains under the same condition, as evidenced by the SYTOX green staining. However, bacterial concentrations increased over time under the asymmetric coculture condition, suggesting that the viability of *Fn* was maintained (**Figure 2D** and **2E**). To confirm if the improved viability was due to the limited oxygen exposure in the apical chamber of the asymmetric coculture system, we performed a control experiment in which *Fn* strains were added to the apical chamber without the differentiated cell monolayer of TIGKs seeded in Transwells. As expected, in the absence of the intact monolayer, a marked reduction in bacterial viability was observed for both strains over time (**Figure S4**). Therefore, our findings underscored the essential role of the intact differentiated TIGK monolayer in generating an oxygen barrier between the aerobic basolateral chamber and the anaerobic apical chamber. Additionally, the asymmetric gas coculture setup offers an advantage over the conventional coculture system with uniform normoxic conditions to simultaneously support the viability of gingival epithelial cells and anaerobic oral bacteria.

### Differential effects of *Fn* challenge on TIGKs under normoxic and asymmetric conditions

Impaired interactions between bacteria and their surrounding epithelium are responsible for bacterial infections (Meyer et al., 1997) (Finlay and Falkow, 1989) (Finlay and Cossart, 1997). To assess the impact of *Fn* on the host cells in the asymmetric coculture system, we examined the changes in the morphology of the TIGK monolayer directly in the Transwell after coculture with three different *Fn* type strains (*Fn* 23726, *Fn* 25586, and *Fn* 10953) or the medium control for 24 h. In the medium control group, the cell monolayer displayed distinct cell junctions, with uniformly sized cells forming a flat, epithelial-like layer. In contrast, the cell monolayer cocultured with *Fn* exhibited irregular morphology with dense bacterial clusters adhering to the surface (indicated by red arrows) (**Figure 3A–3D**). Although the same MOI of 1 was used for all three *Fn* type strains, the formation of dense bacterial clusters was particularly pronounced in strains *Fn* 25586 and 10953, likely due to differences at the sub-species level (Gursoy et al., 2010) (Muchova et al., 2022). Nevertheless, these observations align with previous observations on co-aggregation and adhesive properties of *Fn* during interactions with oral epithelial cells (Han et al., 2000).

**Figure 3:**
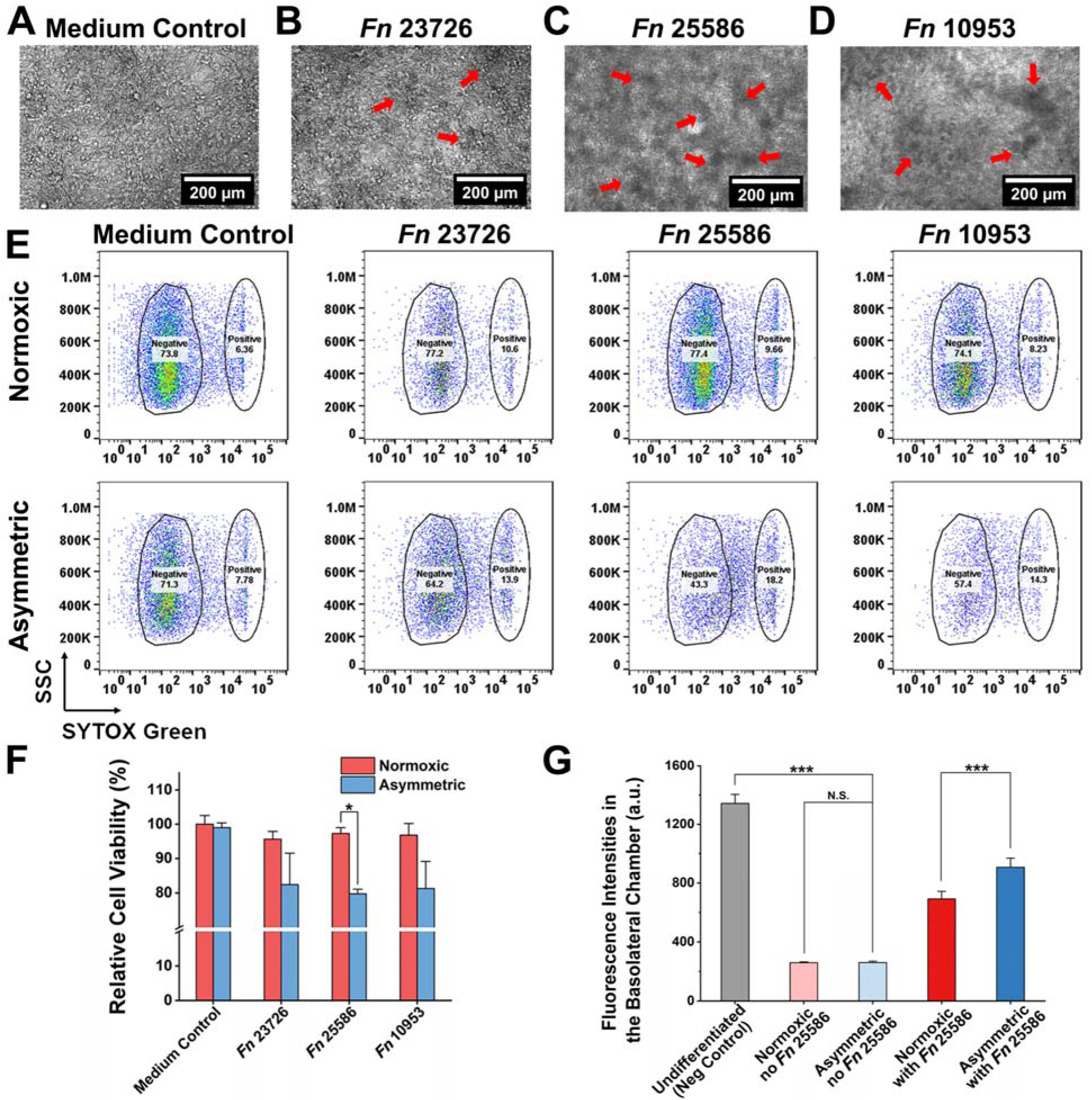
Differential responses of TIGKs challenged by three *Fn* type strains under normoxic and asymmetric gas coculture conditions. Morphological comparison of TIGK monolayers in (A) the medium control (CCM only) and after coculture with (B) *Fn* 23726, (C) *Fn* 25586, and (D) *Fn* 10953 under the asymmetric gas coculture system. The MOI of *Fn* was 1. Red arrows indicate the adherence of bacterial aggregates to the TIGK monolayer. Due to the presence of collagen fibers, collagen coating can obscure fine details of the cells or structures being observed compared to an uncoated surface (Hashimoto et al., 2020). (E) Representative flow cytometry results for TIGKs after coculture with three different *Fn* strains, as assessed by the SYTOX green staining. (F) Quantification of TIGK viability after asymmetric coculture with *Fn* 23726, *Fn* 25586, and *Fn* 10953 based on the flow cytometry results in (E) (*: *p*<0.05, n=2, N=3). (G) Comparison of fluorescence intensities of FITC-dextran in the basolateral chamber at 24 h after adding FITC-dextran to the apical chamber containing TIGK monolayers receiving different treatment conditions. Undifferentiated TIGKs cultured under the normoxic condition served as the negative control due to the lack of an intact monolayer. Differentiated TIGKs with or without *Fn* 25586 challenge under normoxic or asymmetric culture conditions were compared. The background fluorescence of the culture medium was subtracted. The MOI was 1 (N.S.: *p*>0.05, ***: *p*<0.001, n=2, N=3).

We further characterized and compared the adverse effects of three different *Fn* type strains (*Fn* 23726, *Fn* 25586, and *Fn* 10953) on the viability of TIGKs under two different coculture conditions. Compared to the medium control (i.e., no bacterium in the apical chamber), coculture with *Fn* 23726, 25586, and 10953 resulted in ∼20% reductions in TIGK cell viability under the asymmetric coculture system (**Figure 3E** and **3F**). In comparison, there was no noticeable reduction in the viability of TIGKs under normoxic coculture conditions, likely because the viability of *Fn* strains was largely inhibited under the normoxic condition (**Figure 2D** and **2E**). Since *Fn* 25586 exhibited a statistically significant differential impact on the viability of TIGKs between asymmetric and normoxic coculture conditions, we further performed the FITC-dextran diffusion assay, in which the diffusion of FITC-dextran inversely correlates with the integrity of the cell monolayer. It was demonstrated that *Fn* 25586 increased monolayer permeability under asymmetric conditions, with fluorescence intensity in the basolateral chamber increasing by 50% over normoxic conditions (**Figure 3G**). In summary, these results emphasize the importance of the asymmetric coculture condition in demonstrating the severity of *Fn* infection and the impaired viability and barrier function of oral epithelial cells under physiologically relevant conditions.

### Enhanced invasion and pro-inflammatory responses under asymmetric conditions

*Fn* is well known for its adhesion and invasion of oral epithelial cells (Groeger et al., 2022). Adherence to epithelial cells is necessary for colonization, while invasion allows the bacteria to evade immune surveillance and spread to distant organs (Casasanta et al., 2020) (Han et al., 2000) (Meyer et al., 1997) (**Figure 4A**). Based on the above findings on the increased *Fn* viability and severity of *Fn* infection (**Figure 2** and **3**), we hypothesize that the asymmetric coculture conditions may induce elevated adhesion and invasion of *Fn* to oral gingival epithelial cells. To test the hypothesis, we pre-labeled *Fn* with a red fluorescence dye (DiD) and conducted a short 2-h coculture with TIGKs to minimize the impact of bacterial proliferation (**Figure 4B–E** and **Figure S5**). After 2 h, TIGK monolayers were extensively washed to remove free bacteria and stained by Alexa Fluor™ 488 Phalloidin specific for the eukaryotic cytoskeleton. Confocal imaging data suggest that *Fn* was more adherent under asymmetric conditions, particularly for *Fn* 25586 and 10953, indicating strain-dependent variations in invasiveness. To further remove unwashed extracellular bacteria to quantify intracellular bacterial loads, TIGKs were dissociated by trypsin and subjected to a cell-impermeable antibiotic, gentamicin, for 2 h. Subsequently, intracellular bacteria were released upon lysis of TIGKs by a surfactant (Triton X-100), and bacterial counts were quantified by plating serially diluted cell lysates on Columbia blood agar plates. As expected, the same trend was observed for each group’s plating result after coculture. Within 2 h of coculture, *Fn* invasion was significantly higher under asymmetric coculture conditions than under the normoxic condition (2 h, an MOI of 50, **Figure S6**). *Fn* 25586 exhibited greater invasion into TIGKs than *Fn* 23726, further supporting the differential invasiveness of different *Fn* strains (Dabija-Wolter et al., 2009). Although high MOIs, such as ≥50, were typically used for *Fn* coculture with host cells under the normoxic condition in the literature (Liu et al., 2022) (Udayasuryan et al., 2024), we reason that an extremely high MOI may not be physiologically relevant. Therefore, the monolayers of TIGKs were further infected with three different *Fn* strains with an MOI of 1 for 24 h under normoxic or asymmetric coculture conditions. Afterward, extracellular bacteria were eliminated by gentamicin. Intracellular bacterial counts were quantified by plating serially diluted cell lysates on Columbia blood agar plates. Under the asymmetric coculture condition, the intracellular bacteria counts were approximately 10^5^ times higher than those under the normoxic coculture condition, where intracellular bacteria were almost undetectable (**Figure 4F**). This observation was consistent with the increased bacterial proliferation under the asymmetric coculture (**Figure 2**). Furthermore, *Fn* 25586 and 10953 exhibited higher intracellular bacterial loads with the same initial MOI of 1 than *Fn* 23726 under the asymmetric coculture condition (**Figure 4F**). These findings demonstrate that our asymmetric anaerobic coculture system effectively preserves the invasive characteristics of *Fn* in a physiologically relevant environment, which can be severely affected by normoxic conditions.

**Figure 4:**
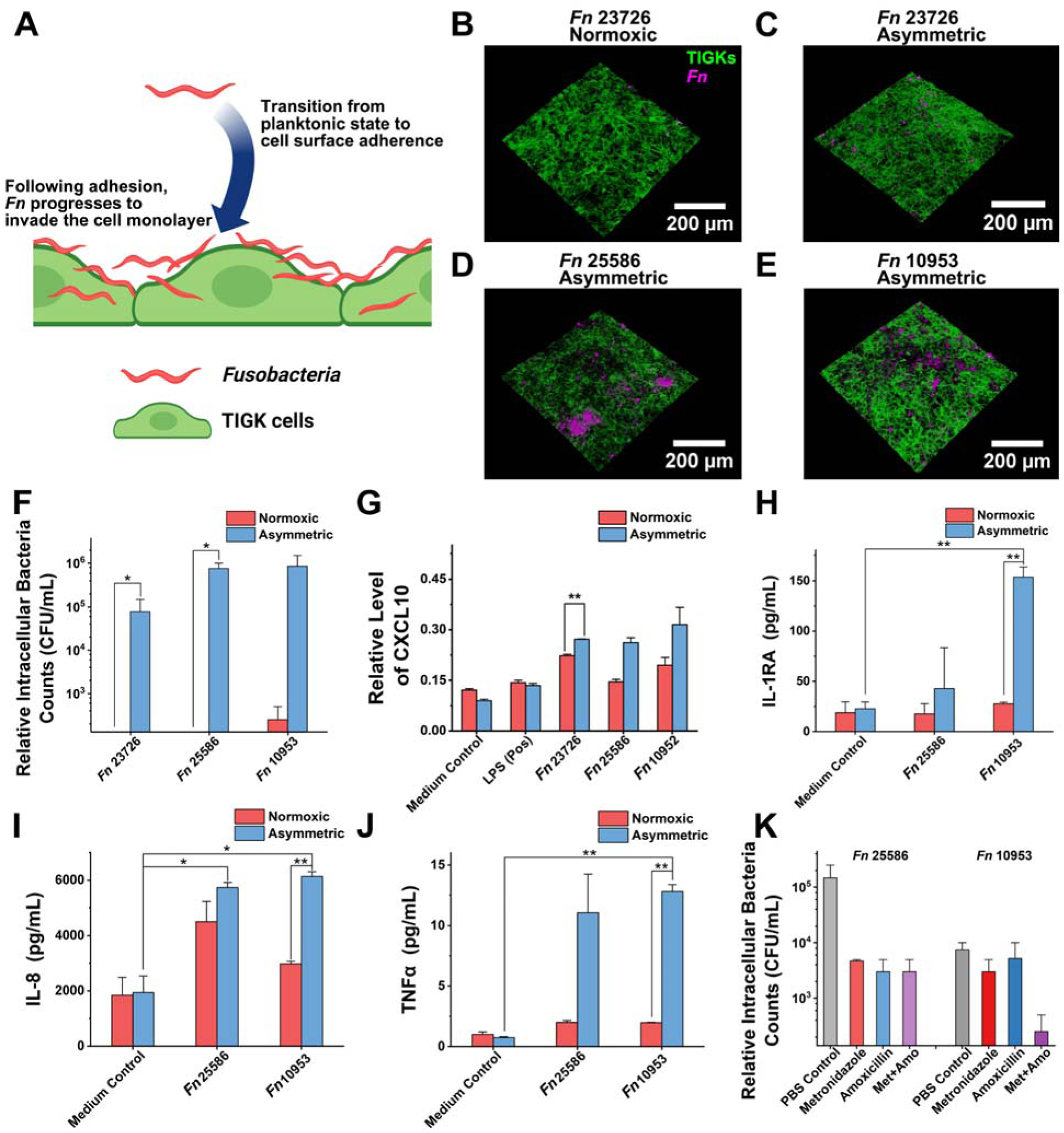
Analysis of *Fn* invasion and its effect on the secretion of proinflammatory factors by cell monolayers. (A) Illustration of *Fn* transitioning from a planktonic state to adherence and invasion to gingival epithelial cells. The figure was created using BioRender. Confocal images showing the interactions between *Fn* and TIGK cell monolayer under normoxic (B) and asymmetric coculture conditions, colocalized with *Fn* strains 23726 (C), 25586 (D), and 10953 (E). *Fn* was prelabeled with a red fluorescence dye DiD, and TIGK cells were post-labeled by Alexa Fluor™ 488 Phalloidin for cytoskeleton. MOI of *Fn* was 50. (F) Differential invasion of three *Fn* strains into the gingival epithelial cells (TIGKs) under normoxic and asymmetric coculture conditions for 24 h. The MOI of *Fn* was 1 (*: 0.01<*p*<0.05; n=2, N=3). (G) Enzyme-linked immunosorbent assay result of CXCL10 (IP-10) expression levels after infection with different *Fn* strains. The value was measured by assessing absorbance at 450 nm, with a reference at 570 nm (n=3, N=3). (H)-(J) Luminex outcome of the difference in expression levels of different pro-inflammatory factors after infection with different *Fn* strains (*: 0.01<*p*<0.05, **: *p*<0.01; n=1, N=3). (K) Intracellular bacterial counts of *Fn* 25586 and *Fn* 10953 after treatment with amoxicillin, metronidazole, and their combination (n=2, N=3).

Besides serving as a physical barrier, the oral gingival epithelium also provides early immune surveillance to alert the presence of pathogenic bacteria. The direct contact between bacteria and the mucosal surface elicits the secretion of different immune cytokines and chemokines from epithelial cells (Neurath and Kesting, 2024) (Wright et al., 2011) (Okada and Murakami, 1998) (Kagnoff and Eckmann, 1997). Considering the higher invasiveness displayed by *Fn* 25586 and *Fn* 10953 over *Fn* 23726 (**Figure 3** and **Figure 4F**), we focused on *Fn* 25586 and *Fn* 10953 to evaluate the differential inflammatory responses from TIGKs under normoxic and asymmetric conditions. CXCL10 (IP-10), a key chemokine in the CXC chemokine family, plays a critical role in recruiting different immune cell types and is heavily involved in inflammation and immune regulation. Given the established link between elevated CXCL10 levels and oral bacterial infections of host cells (Jang et al., 2017) (Groeger et al., 2022), we quantified the induction of CXCL10 using an enzyme-linked immunosorbent assay. Our results indicated that CXCL10 levels in the bacterial challenge group increased more than twofold compared to the baseline level of the unchallenged group under the asymmetric gas coculture conditions (**Figure 4G**). In contrast, under normoxic conditions, the increase in CXCL10 levels in the bacterial challenge group was less than one-fold compared to the unchallenged group. Furthermore, Luminex analysis of additional pro-inflammatory markers revealed significant upregulation of IL-6, IL-8, TNF-α, and IL-1RA (**Figure 4H–4J** and **Figure S7**), consistent with previous studies (Casasanta et al., 2020) (Koido et al., 2022) (Sheikhi et al., 2000). Of note, the expression of IL-8, a proinflammatory chemokine, has been suggested to be an important regulatory mechanism to induce neutrophil migration into the gingival sulcus (Bickel, 1993). Taken together, these results highlight the ability of the asymmetric gas coculture system to effectively mimic the proinflammatory responses and cytokine/chemokine secretion typically observed during oral infections, offering an advantage over conventional coculture models for studying bacteria-host interactions in a physiologically relevant environment.

### Antibiotic susceptibility of intracellular *Fn* under asymmetric conditions

To demonstrate the translational potential for applied research, we explored the utility of the asymmetric coculture system for evaluating antibiotic treatment of bacterial infection. The invasion of *Fn* into host cells may contribute to increased resistance to antibiotics, leading to secondary infections (Mountcastle et al., 2020). To assess the efficacy of commonly used antibiotics against intracellular *Fn*, in addition to gentamicin, we selected metronidazole and amoxicillin, two widely prescribed treatments for oral inflammation (Brennan and Garrett, 2019). We first demonstrated that extracellular *Fn* in the apical chamber of the asymmetric coculture was effectively inhibited by individual antibiotics, reducing bacterial counts by over three orders of magnitude compared to the untreated group (**Figure S8**). We then treated TIGKs with each antibiotic to investigate their effects on intracellular bacteria. Metronidazole and amoxicillin individually exhibited certain inhibitory efficacy against intracellular *Fn* 25586, reducing bacterial load by an order of magnitude, while their combination did not further reduce the bacterial loads (**Figure 4K**). In comparison, metronidazole and amoxicillin inhibited intracellular *Fn* 10953, respectively, and their combined use significantly decreased intracellular bacterial load, reducing it to as low as 1% of the original level, which aligns with previous reports (Anyiam et al., 2024) (Dabija-Wolter et al., 2018) (Feres et al., 2021). These findings highlight the potential of our asymmetric oral coculture platform as a valuable tool for screening antibiotic susceptibility against intracellular oral pathogenic anaerobes, facilitating the development of targeted therapeutic drugs.

## Discussion

This study highlights the critical need for physiologically relevant *in vitro* models to study microbial-host interactions, particularly for anaerobic pathogens like *Fusobacterium nucleatum* (*Fn*). Existing coculture systems, while instrumental, fail to mimic the oxygen gradients inherent to the oral environment. The asymmetric gas coculture system developed here addresses these limitations by creating a controlled oxygen gradient that preserves bacterial viability and epithelial cell functionality. This innovation is particularly significant for studying anaerobes, whose growth and pathogenic potential are often compromised in traditional setups. The system’s ability to maintain the integrity of both host and microbial components over extended periods provides a robust platform for investigating complex interactions, including bacterial adhesion, invasion, and immune modulation.

Regarding microbial pathogenesis and host responses, this work demonstrates that the asymmetric coculture model not only sustains bacterial viability but also recapitulates key pathogenic features of *Fn*, such as enhanced invasion and intracellular persistence. These capabilities are pivotal for studying how *Fn* contributes to periodontal disease and its potential systemic implications, including its role in cancers. Moreover, the increased secretion of pro-inflammatory cytokines like CXCL10 and IL-8 under asymmetric conditions underscores the model’s capacity to mimic innate immune responses observed *in vivo*. By enabling the study of pathogen virulence and host immune dynamics, this system offers a nuanced view of the molecular and cellular mechanisms underpinning microbial pathogenesis and host defense.

Beyond advancing fundamental understanding, the asymmetric gas coculture system has significant implications for drug discovery and development. The differential antibiotic susceptibility observed for intracellular *Fn* highlights the importance of testing therapeutics under conditions that closely simulate the physiological microenvironment. The system’s ability to differentiate the effects of antibiotics on extracellular versus intracellular bacteria presents a valuable tool for evaluating targeted therapies. Furthermore, by maintaining the physiological relevance of bacterial and host interactions, this model could streamline the development of novel therapeutics for oral infections and related systemic diseases, bridging the gap between *in vitro* studies and clinical applications.

Future work could focus on improving the system’s capabilities by incorporating a flow-based setup that continuously supplies fresh medium to both bacterial and host compartments (Makkar et al., 2023). Such an enhancement would allow for extended investigations of microbial-host dynamics over longer time frames, providing insights into chronic interactions and adaptations. Moreover, evaluating the system with a broader range of cell types, such as immune cells or fibroblasts, and oral commensals or pathobionts, would enable the exploration of differential host responses to various microbial species (Adelfio et al., 2023) (Welch et al., 2016). These advancements could reveal key mechanisms driving oral health and disease and further establish the platform’s utility for personalized medicine and therapeutic screening.

## Acknowledgments

This work was supported by National Science Foundation CAREER (2238972) and National Institute of Dental and Craniofacial Research awards (R03DE031329 and R01DE030943). We thank the Translational Tissue Modeling Laboratory and the Microscopy Core at the University of Michigan, Ann Arbor, for their valuable support. We are especially grateful to Binyamin Jacobovitz from the Microscopy Core for his assistance with confocal microscopy equipment and training. We also extend our thanks to Michael K. Dame, Jack Morgan, and Dominic Tigani from the Translational Tissue Modeling Laboratory for their expert training and insightful suggestions on cell monolayer preparation techniques. The Translational Tissue Modeling Laboratory is supported by the University of Michigan (Center for Gastrointestinal Research, Office of the Dean, Comprehensive Cancer Center, and the Departments of Pathology, Pharmacology, and Internal Medicine) with additional funding from the Endowment for Basic Sciences.

## Data Availability

All data generated during this study are available within the paper.

