## Supporting Information for "Physiologically Relevant Coculture Model for Oral Microbial-Host Interactions"

**Materials and Methods**

*Chemicals*

Unless otherwise noted, all chemicals and cell culture broth were purchased from Fisher Scientific International Inc. (Cambridge, MA, USA) and were commercially available of the highest purity or analytical grade. Ultrapure water is provided by the laboratory's Mili-Q water purification system (MilliporeSigma, Darmstadt, Germany).

*Bacterial Strains and Growth Conditions*

*Fusobacterium nucleatum strains* (*Fn*, ATCC 23726, 25586, and 10953) were purchased from the American Type Culture Collection (Manassas, VA, USA). All strains were cultured in liquid Columbia broth (CB) or on CB agar plates containing 5% defibrinated sheep blood (Hemostat laboratories, Dixon, CA, USA) and incubated at 37°C in an anaerobic chamber (Type A, Coy Laboratories, Grass Lake, MI, USA) containing 5% H_2_, 10% CO_2_, 85% N_2_ (Cryogenic Gases, Ypsilanti, MI).

*Cell Strains and Culture Conditions*

The cell line used in this manuscript is the hTERT TIGKs cell line (CRL-3397, ATCC). The base medium used was Dermal Cell Basal Medium (PCS-200-030, ATCC) for culture, to which the components of the Keratinocyte Growth Kit (PCS-200-040, ATCC) were added. The culture atmosphere was 95% Air + 5% CO_2_, and the temperature was 37°C.

*SYTOX Green Assay*

SYTOX green (S7020, ThermoFisher, Waltham, MA, USA) was added to *Fn* or corresponding cell monolayer apical chamber at a working concentration of 5 μM in phosphate buffer saline (PBS). Fluorescence intensity was then acquired after incubation at room temperature for 10 min. SYTOX green was excited at a wavelength of 488 nm and collected through a bandpass filter from 500-550 nm. Transmission images were acquired through a differential interference contrast setting. All the images were acquired under the same image acquisition setting and analyzed by *ImageJ*.

*Transwell collagen coating*

All Transwell inserts (76313-906, VWR Inc, Radnor, PA, USA) were precoated with human collagen type IV (C5533, MilliporeSigma) before seeding cells. Specifically, collagen IV was formulated to 1 mg/mL stock solution dissolved in 100 mM acetic acid solution. The collagen stock solution was diluted to 33 μg/mL with ice-cold ultrapure water and mixed well, and 100 μL of diluted solution was aliquoted into each apical chamber of the Transwell inserts. The entire cell plate was incubated at 4°C for 30 min and then placed in a 37°C cell culture incubator for 4 h.

*2D Monolayers*

2D monolayers were seeded at a density of 200,000 cells per 100 μL in each Transwell insert and incubated for 20 h in the same culture medium used for cell maintenance. Simultaneously, 600 μL of the same medium was added to the basolateral chamber of the Transwell. The following day, the culture medium was replaced with Dulbecco's Modified Eagle's Medium (11965092, ThermoFisher) supplemented with 10% fetal bovine serum (16140071, ThermoFisher) and 100 U/mL Penicillin-Streptomycin (15140122, ThermoFisher). Differentiation was then allowed to proceed for 5 days.

*Asymmetrical Coculture*

One day before the experiment, *Fn* was inoculated at a 1:100 ratio into 1 mL of cell-compatible medium (CCM; see **Supplementary Table S1** for the specific ingredient list) and grown until the optical density at 600 nm (OD600) reached ~0.4. The medium in the Transwell inserts was replaced with DMEM supplemented with 10% FBS. The inserts were then transferred into an asymmetrical gas system consisting of a gas-permeable plate (8704000, Coy Laboratories) placed inside an anaerobic chamber maintained at 37°C. A gas mixture of 10% oxygen, 5% CO₂, and 85% nitrogen (Cryogenic Gases) was supplied to the basolateral side at a flow rate of 0.5 SCFH. After 3 h of purge, CCM or the *Fn* solution with a multiplicity of infection (MOI)=1 was used to replace the apical chamber medium. To minimize the impact of hypoxia due to limited diffusion caused by the monolayer’s oxygen consumption, the system was placed atop a magnetic stirrer, generating an oxygen gradient in the basolateral chamber. This setup was maintained at 37°C for 24 h.

*Fluorescein Isothiocyanate (FITC)-dextran Permeability Test*

FITC-dextran 4kD (46944, MilliporeSigma) was prepared as a 50 mg/mL working solution. To each apical chamber, 4 μL of the working solution was added to achieve a final 1 mg/mL concentration. The Transwells were then wrapped in aluminum foil to protect them from light and incubated at 37°C in a cell culture incubator for 2 h. After incubation, 200 μL of the basolateral solution was aspirated and transferred to a black flat bottom 96-well Corning fluorescent plate. Fluorescence intensity was measured with 475 nm excitation and 525 nm emission using a 515 nm filter.

*Laser Confocal Microscopy of TIGK monolayers coculture with different Fn Strains*

TIGK cells were precultured using the previously described method until day four to establish a mature cell monolayer. *Fn* was pre-labeled with DiD dye (DiIC18(5); 1,1′-dioctadecyl-3,3,3′,3′- tetramethylindodicarbocyanine, 4-chlorobenzenesulfonate salt) at a final concentration of 10 μM and incubated at 37°C for 30 minutes. The stained *Fn* was then added to the apical chamber containing the cell monolayer at an MOI of 50 and incubated for 2 h in complete darkness. After coculture, cell monolayers were carefully removed and washed three times with PBS under light-proof conditions. The cell monolayers were then fixed with 4% paraformaldehyde solution at 4°C for 30 minutes, followed by three additional PBS washes to eliminate residual paraformaldehyde and prevent interference with subsequent steps. The fixed cell monolayer was carefully cut from the Transwell using a surgical blade and incubated with Alexa Fluor 488 Phalloidin (A12379, Invitrogen), diluted 1:100, at room temperature for 1 h in the dark. The monolayer was washed four times with PBS, each lasting 5 minutes, to remove any excess phalloidin dye. Following the washes, the sample was placed on a slide and mounted using EverBrite Hardset Mounting Medium (23001, Biotium, Fremont, CA, USA). A coverslip was applied, and the entire sample was stored at room temperature in the dark overnight. Fluorescence images were captured using a Nikon A1 inverted confocal microscope with excitation lasers (640 nm for DiD and 488 nm for Alexa Fluor 488 Phalloidin). The Z-stack function was utilized to acquire three-dimensional images of cell monolayers.

*Human CXCL10 ELISA Assay*

All apical chamber solutions were analyzed directly by Human CXCL10/IP-10 DuoSet ELISA kit (DY266-05, R&D Systems, Minneapolis, MN, USA). All operations were performed according to the kit protocol instructions. 100 ng/mL lipopolysaccharide (LPS) was applied as the positive control to stimulate the secretion of CXCL10.

*Luminex Human Cytokine Level Assay*

For sample preparation, the apical chamber solution was centrifuged at 3000 x g three times at 4°C, with the supernatant collected after each spin to remove cellular debris. The resulting cell culture supernatant was aliquoted into pyrogen-free 0.65 mL snap-top tubes and stored in dry ice. The Human Cytokine Proinflammatory Focused 15-Plex Discovery Assay was conducted by Eve Technologies Corporation (Calgary, AB, Canada). Multiplex analysis was performed using the Luminex™ 200 system (Luminex, Austin, TX, USA). According to the manufacturer's protocol, fifteen markers were measured simultaneously in samples using the Human-Focused 15-Plex Discovery Assay from Eve Technologies (MilliporeSigma, Burlington, MA, USA). 15 consists of GM-CSF, IFNγ, IL-1β, IL-1Ra, IL-2, IL-4, IL-5, IL-6, IL-8, IL-10, IL-12p40, IL-12p70, IL -13, MCP-1 and TNF-α. Detection sensitivities for these markers ranged from 0.14 – 5.39 pg/mL for the 13-plex. Sensitivity values for individual analytes are provided in the MilliporeSigma MILLIPLEX® MAP protocol.

*Bacteria level analysis*

After the coculture was completed, the apical chamber solution was taken to a serial dilution. Specifically, 90 µL of PBS was added to each well of a 96-well plate, followed by 10 µL of the apical chamber solution to the first well of each row. After thorough mixing, 10 µL was transferred to the second well, and this process was repeated to achieve a tenfold serial dilution across the plate. Bacterial suspensions from each group were then plated on CB blood agar plates, allowing for the determination of the original bacterial count through back-calculation.

For intracellular bacterial counting, following the coculture phase, cells in the Transwell inserts were washed with PBS and digested with 0.25% trypsin for 4 min. The digested cells were then resuspended in DMEM containing 50 µg/mL of gentamicin and incubated at 37°C for 2 h to eliminate any remaining extracellular bacteria. After incubation, the cells were washed twice with PBS and lysed using a solution of 0.1% Triton X-100 and 0.01% SDS in PBS (Li et al., 2023). The lysates were serially diluted as previously described and plated on CB blood agar plates to quantify the intracellular bacteria. For all other antibiotic treatment groups, the antibiotic concentration was adjusted to 200 µg/mL metronidazole, 200 µg/mL amoxicillin, and the combination of 100 µg/mL metronidazole + 100 µg/mL amoxicillin (Robinson et al., 2024).

*Statistical Analysis*

All statistical analyses were performed using GraphPad Prism 9 (San Diego, CA, USA). Data were analyzed using the student t-test, a one-way or two-way analysis of variance (ANOVA), followed by the *Bonferroni* test for statistical significance.

**Supporting Tables and Figures**

**Table S1. Cell-compatible media (CCM) ingredient list**

| Reagents | Working Concentrations |
| --- | --- |
| Mineral Solutions |  |
| Dipotassium phosphate (KH_2_PO4) | 0.51 mM |
| Potassium Diphosphate (K_2_HPO4) | 2.94 mM |
| Potassium Chloride (KCl) | 5.37 mM |
| Magnesium sulfate heptahydrate (MgSO_4_·7H_2_O) | 0.41 mM |
| Calcium chloride dihydrate (CaCl_2_·2H_2_O) | 1.26 mM |
| Magnesium dichloride hexahydrate (MgCl_2_·6H_2_O) | 0.49 mM |
| Ammonium Sulfate [(NH_4_)_2_SO_4_] | 3.4 mM |
| Sodium Chloride | 8 g/L |
| Vitamin Mix |  |
| Biotin | 8.19 μM |
| Folic Acid | 4.53 μM |
| Pyridoxine HCl | 48.63 μM |
| Thiamine HCl | 14.82 μM |
| Riboflavin | 13.26 μM |
| Nicotinic acid | 40.61 μM |
| D-Pantothenic acid hemicalcium salt | 20.98 μM |
| Vitamin B12 | 7.37 μM |
| *p*-Aminobenzoic acid (PABA) | 36.46 μM |
| α-Lipoic acid (thioctic acid) | 24.23 μM |
| Trace Metals |  |
| EDTA | 17.1 μM |
| FeSO_4_·7H_2_O | 3.60 μM |
| ZnSO_4_·7H_2_O | 6.26 μM |
| CuSO_4_·7H_2_O | 350 nM |
| CoCl_2_·6H_2_O | 7.57 μM |
| MnSO_4_·H_2_O | 29.58 μM |
| NiCl_2_·6H_2_O | 2.95 μM |
| Glucose | 0.9 g/L |
| Yeast Extract | 1.25 g/L |
| Tryptone | 5 g/L |
| Haemin | 2 g/L |
| Bromocresol purple | 1 g/L |
| Sodium bicarbonate | 0.338 g/L |

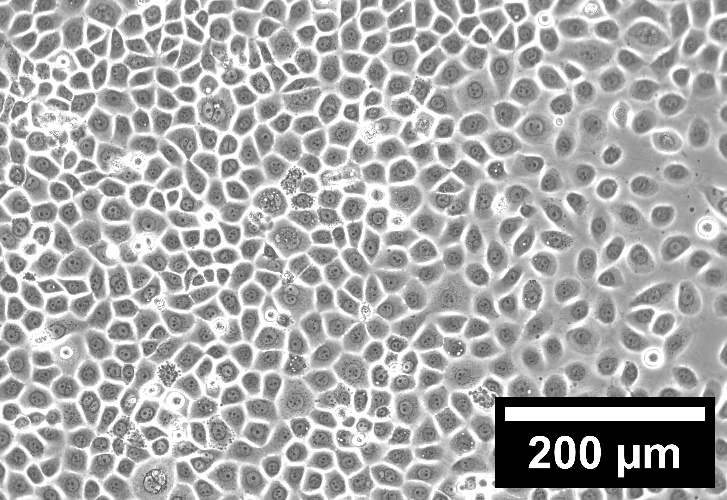

**Figure S1:** Cell morphology of cultured hTERT Telomerase Immortalized Gingival Keratinocytes (TIGKs) before seeding into Transwell inserts in well plates.

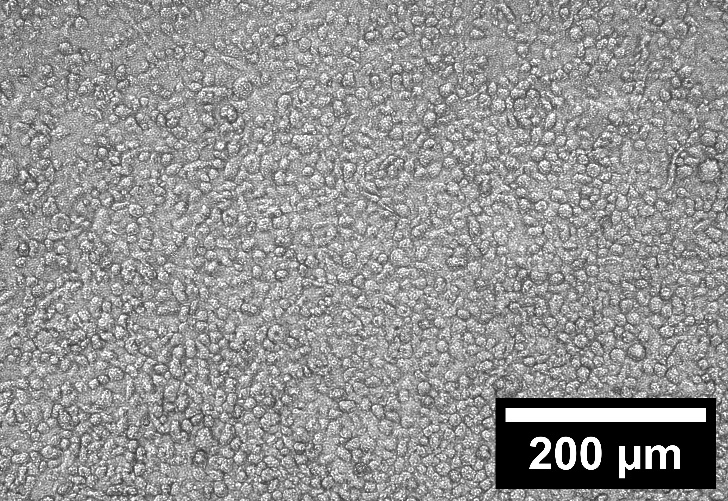

**Figure S2:** Cell morphology of hTERT-TIGK cell monolayer in collagen IV-coated Transwell inserts after 4 d of differentiation within DMEM containing calcium ions. TIGKs form a tight monolayer on the bottom membrane of the Transwell.

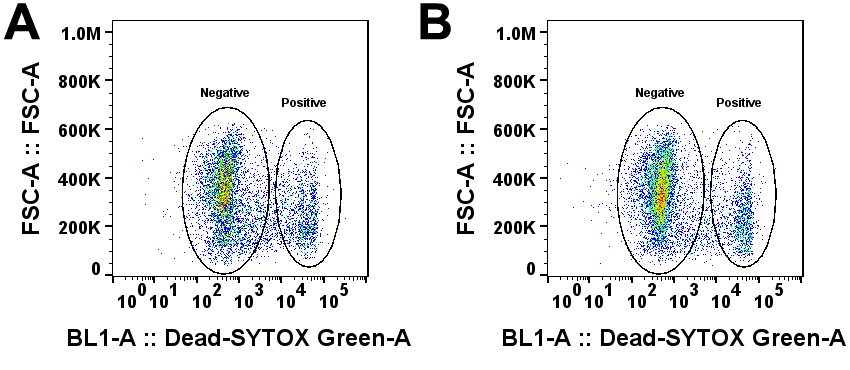

**Figure S3:** Representative flow cytometry results for the viability of TIGKs monolayer cultured under (A) normoxic conditions and (B) asymmetric conditions without bacterial challenge.

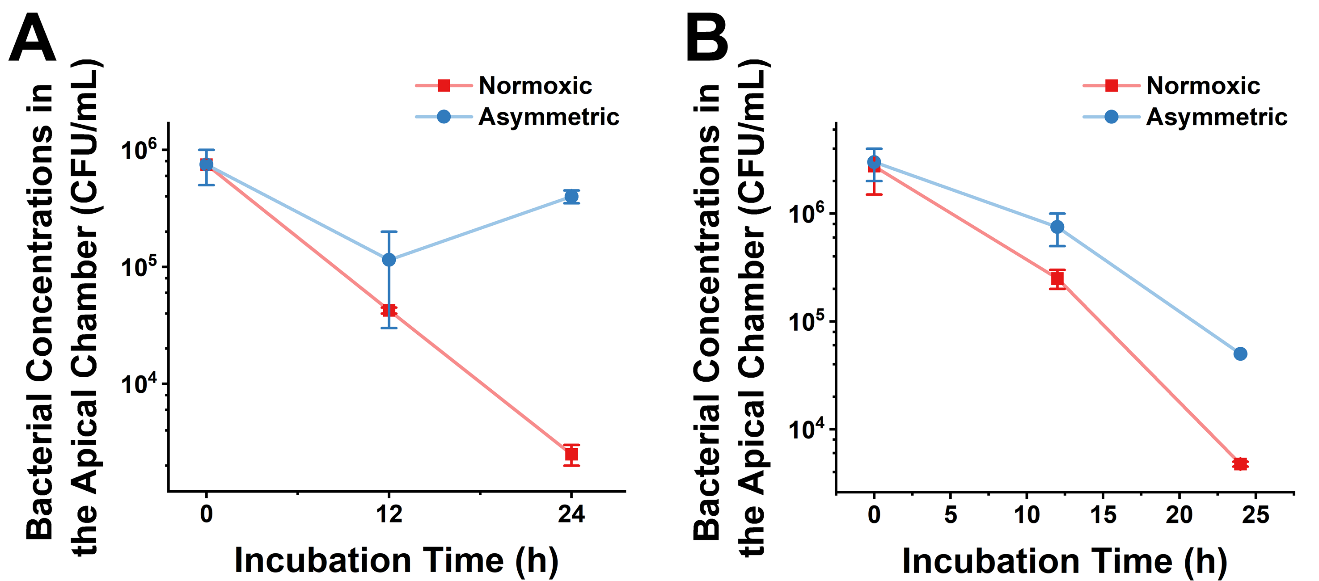

**Figure S4:** (A) Comparison of Fn 23726 concentration changes over 0, 12, and 24 h in the apical chambers without TIGK cell monolayer under normoxic and asymmetric coculture conditions. (B) Comparison of Fn 25586 concentration changes over 0, 12, and 24 h in the apical chambers without TIGK cell monolayer under normoxic and asymmetric coculture conditions (n=2, N=3).

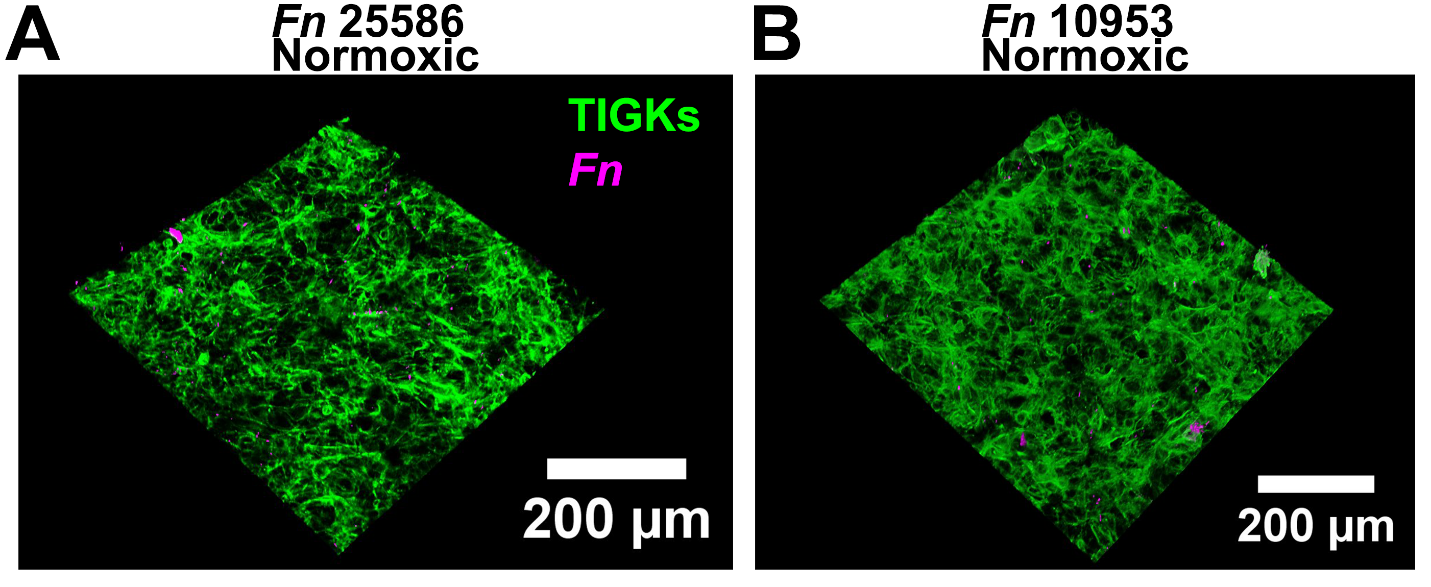

**Figure S5:** Confocal images showing minimal adhesion and invasion of (A) Fn 25586 and (B) Fn 10953 in the TIGK monolayer under normoxic coculture conditions.

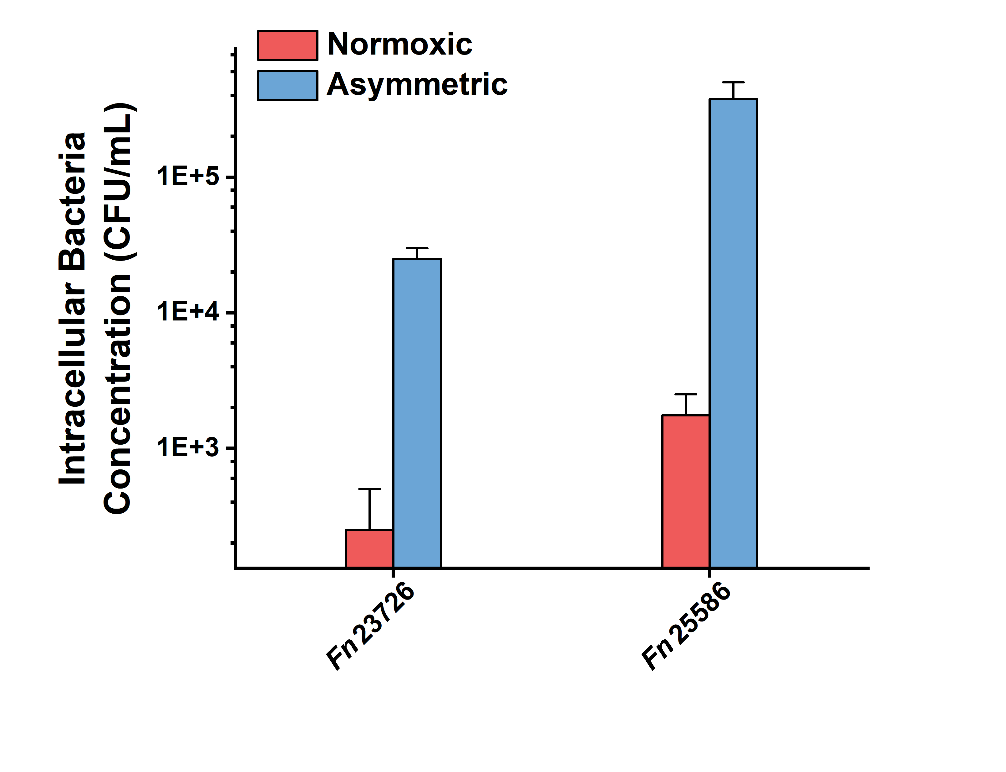

**Figure S6:** Analysis of Fn 23726 and 25586 intracellular bacterial concentrations after 2 h of normoxic and asymmetric coculture. Notably, during the 2-h cell invasion under normoxic conditions, the activity of Fn was not fully inhibited due to its brief exposure to oxygen. As a result, under the same starting MOI, the intracellular CFU count was higher than the 24-h normoxic counterparts (n=2, N=3).

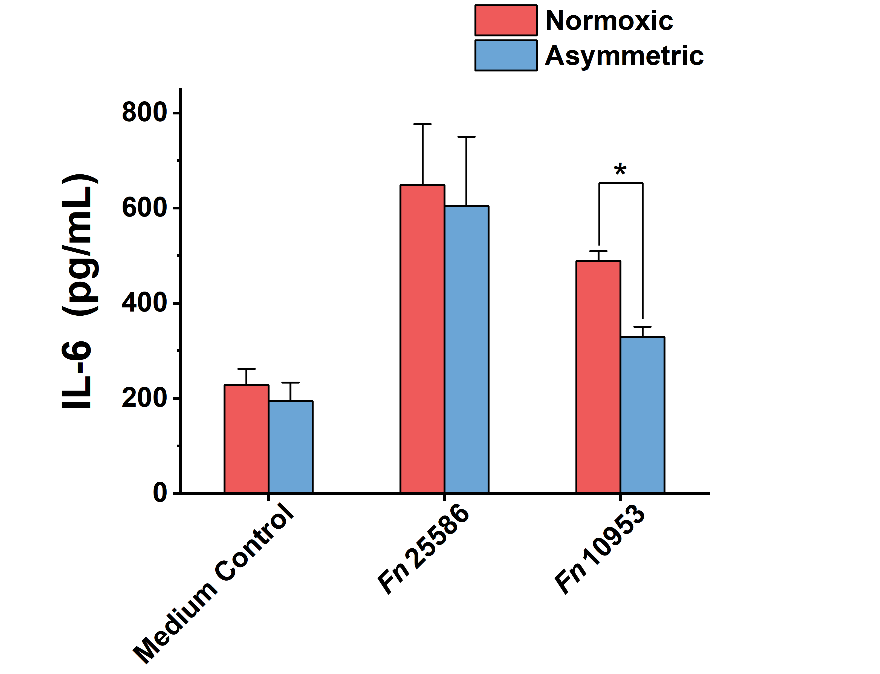

**Figure S7:** Luminex analysis showing differences in IL-6 expression levels in TIGK cell monolayers following infection with various Fn strains (*: 0.01<p<0.05, n=1, N=3).

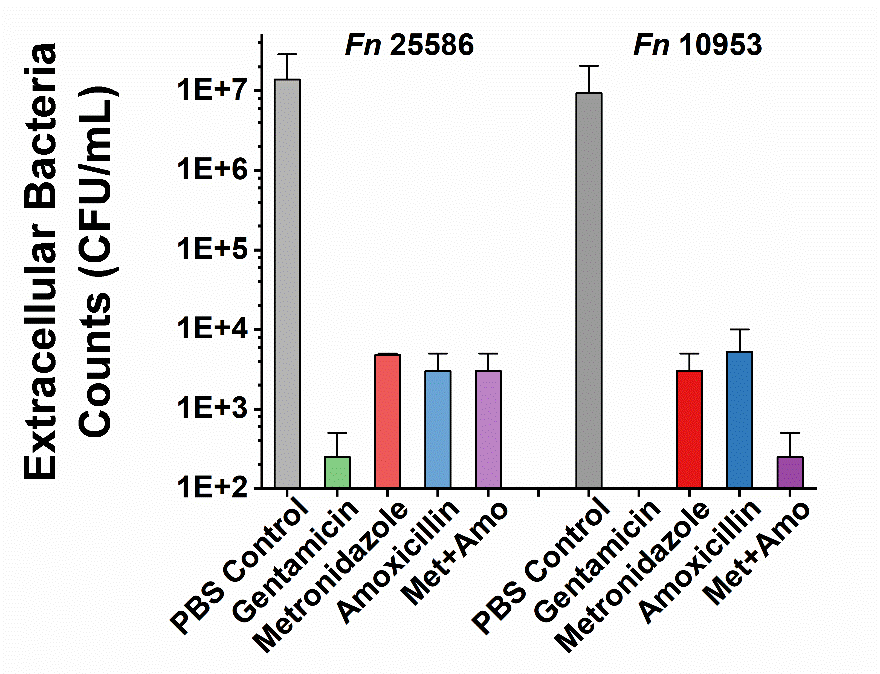

**Figure S8:** Extracellular concentrations of Fn 25586 and 10953 after treatment with PBS and the same concentrations of gentamicin, amoxicillin, metronidazole, and their combination were used for evaluating intracellular bacteria (n=2, N=3).
